# Multi-context blind source separation by error-gated Hebbian rule

**DOI:** 10.1101/441618

**Authors:** Takuya Isomura, Taro Toyoizumi

## Abstract

Animals need to adjust their inferences according to the context they are in. This is required for the multi-context blind source separation (BSS) task, where an agent needs to infer hidden sources from their context-dependent mixtures. The agent is expected to invert this mixing process for all contexts. Here, we show that a neural network that implements the *error-gated Hebbian rule* (EGHR) with sufficiently redundant sensory inputs can successfully learn this task. After training, the network can perform the multi-context BSS without further updating synapses, by retaining memories of all experienced contexts. Finally, if there is a common feature shared across contexts, the EGHR can extract it and generalize the task to even inexperienced contexts. This demonstrates an attractive use of the EGHR for dimensionality reduction by extracting common sources across contexts. The results highlight the utility of the EGHR as a model for perceptual adaptation in animals.

## Introduction

Inference of the causes of a sensory input is one of the most essential abilities of animals (*Helmholtz, 1925; Knill, Pouget, 2004; DiCarlo et al, 2012*) –– a famous example is the cocktail party effect, i.e., the ability of a partygoer to distinguish a particular speaker’s voice against a background of crowd noise (*Brown et al., 2001; Mesgarani, Chang, 2012*). This ability has been modelled by blind source separation (BSS) algorithms (*Cichocki et al., 2009; Comon, Jutten, 2010*), by considering that several hidden sources (speakers) independently generate signal trains (voices), while an agent receives mixtures of signals as sensory inputs. A neural network, possibly inside the brain, can invert this mixing process and separate these sensory inputs into hidden sources using a BSS algorithm. Independent component analysis (ICA), achieves BSS by minimizing the dependency between output units (*Comon, 1994; Comon, Jutten, 2010*). Numerous ICA algorithms have been proposed for both rate-coding (*Bell, Sejnowski, 1995; Bell, Sejnowski, 1997; Amari et al., 1996; Hyvarinen, Oja, 1997*) and spiking neural networks (*Savin et al., 2010*).

Previously, we developed a biologically plausible ICA algorithm, referred to as the *error-gated Hebbian rule (EGHR)* (*Isomura, Toyoizumi, 2016*). This learning rule can robustly estimate the hidden sources that generate sensory data without supervised signals. Importantly, it can reliably perform ICA in undercomplete conditions (*Lee, Girolami et al., 2000*), where the number of inputs is greater than that of outputs. A simple extension of the EGHR can separate sources while removing noise within a single-layer neural network (*Isomura, Toyoizumi, 2018*), by simultaneously performing principal component analysis (PCA) (*Pearson, 1901; Oja, 1989*) and ICA. The EGHR is expressed as a product of pre-and post-synaptic neuronal activities and a third modulatory factor, each of which can be computed locally (i.e., local learning rule (*Lee, Girolami et al., 2000*)). In this sense, the EGHR is more biologically plausible than non-local engineering ICA algorithms (*Bell, Sejnowski, 1995; Bell, Sejnowski, 1997; Amari et al., 1996*). Because of these desirable properties, the EGHR is considered as a candidate mechanism for neurobiological BSS (*Kuśmierz et al., 2017; Avitan, Goodhill, 2018*), as well as a next-generation neuromorphic implementation (*Neftci, 2018*) for energy efficient BSS.

The optimal inference and behavior often depend on context. Indeed, our perception and decisions reflect this context-dependence, i.e., cognitive flexibility (*Dajani, Uddin, 2015*). Studies in primates have suggested that a contextual-cue-dependent dynamic process in the prefrontal cortex controls this behavior (*Dehaene, Changeux, 1991; Gilbert, Sigman, 2007; Mante et al., 2013*), and several computational studies have modeled it (*Mante et al., 2013; Song et al., 2016; Song et al., 2017; Miconi, 2017; Chaisangmongkon et al., 2017*). Likewise, context dependence of auditory perceptual inference has been modeled (*Ahrens et al., 2008*). In addition to experimental evidence, recent progress in machine learning has also addressed this multi-context problem, in an attempt to create artificial general intelligence (*Yu et al., 2010; Kirkpatrick et al., 2017; Zenke et al., 2017*). By implementing (task-specific) synaptic consolidation, a neural network can learn a new environment, while retaining past memories, by protecting synaptic strengths that are important to memorizing past environments. Those findings indicate the importance of multi-context processes for cognitive flexibility.

Unlike the above-mentioned tasks, BSS in several different contexts has some difficulty. Conventional ICA algorithms assume the same number of input and output neurons (*Bell, Sejnowski, 1995; Bell, Sejnowski, 1997; Amari et al., 1996, Foldiak, 1990; Linsker, 1997*) and, thus, cannot straightforwardly perform a multi-context BSS. After learning, the synaptic strength matrix of these algorithms converges to the inverse of the mixing matrix of the current context (or its permutation or sign-flip), which is generally different from that in the previous context. Hence, when the network subsequently encounters a previously learnt context, it needs to relearn the synaptic strengths from the very beginning. More involved engineering ICA algorithms, such as the nonholonomic ICA algorithm (*Amari et al., 2000*) and the ICA mixture algorithm (*Lee, Lewicki et al., 2000; Hirayama et al., 2015*), are expected to perform the multi-context BSS. However, we argue that EGHR has the advantage of being a biologically plausible local learning rule.

Here we show that the EGHR can perform multi-context BSS when a neural network receives redundant sensory inputs. It can retain memories of previously experienced contexts and process the BSS right after contextual switching to a previously learnt context. This suggests that the EGHR can also be used as a powerful data compression method (*Cunningham, Ghahramani, 2015*), since it extracts hidden sources shared across contexts, despite the proportional increase in data dimensions to the number of contexts. Moreover, when a common feature is shared across contexts, the EGHR can extract it to perform BSS, while filtering out features that vary among contexts. Once the learning is achieved, the network can perform BSS even in an inexperienced context, indicating some generalization capability or transfer learning. We demonstrate that the EGHR with sufficiently redundant sensory inputs learns to distinguish birdsongs from their superpositions and retains this ability even after learning different sets of birdsongs. The rule finds a general representation that is capable of separating an unheard set of birdsongs. Finally, possible neurobiological implementations of the EGHR are discussed.

## Results

### Error-gated Hebbian rule (EGHR)

In a BSS task, several hidden sources (*s*) independently generate signal traces, while our agent receives their mixtures as sensory inputs (*x*). In this study, we consider a multi-context BSS task, in which a set of contexts with different mixing weights is used. Sensory inputs are randomly generated from one of these contexts for a period of time. Let *k* (= 1,…, *C*) be an index of context. Our experimental setup consists of an *N_s_*-dimensional vector of hidden Sources 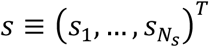 whose elements *s_i_* independently follow a non-Gaussian, distribution *p*(*s_i_*), an *N_x_*-dimensional vector of sensory inputs 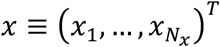,, and an *N_u_*-dimensional vector of neural outputs 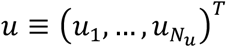 (Fig. 1). The sensory inputs in the *k*-th condition are generated by transforming the hidden sources, i.e., the so-called generative process:

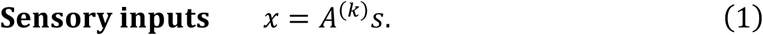

**Figure 1.**
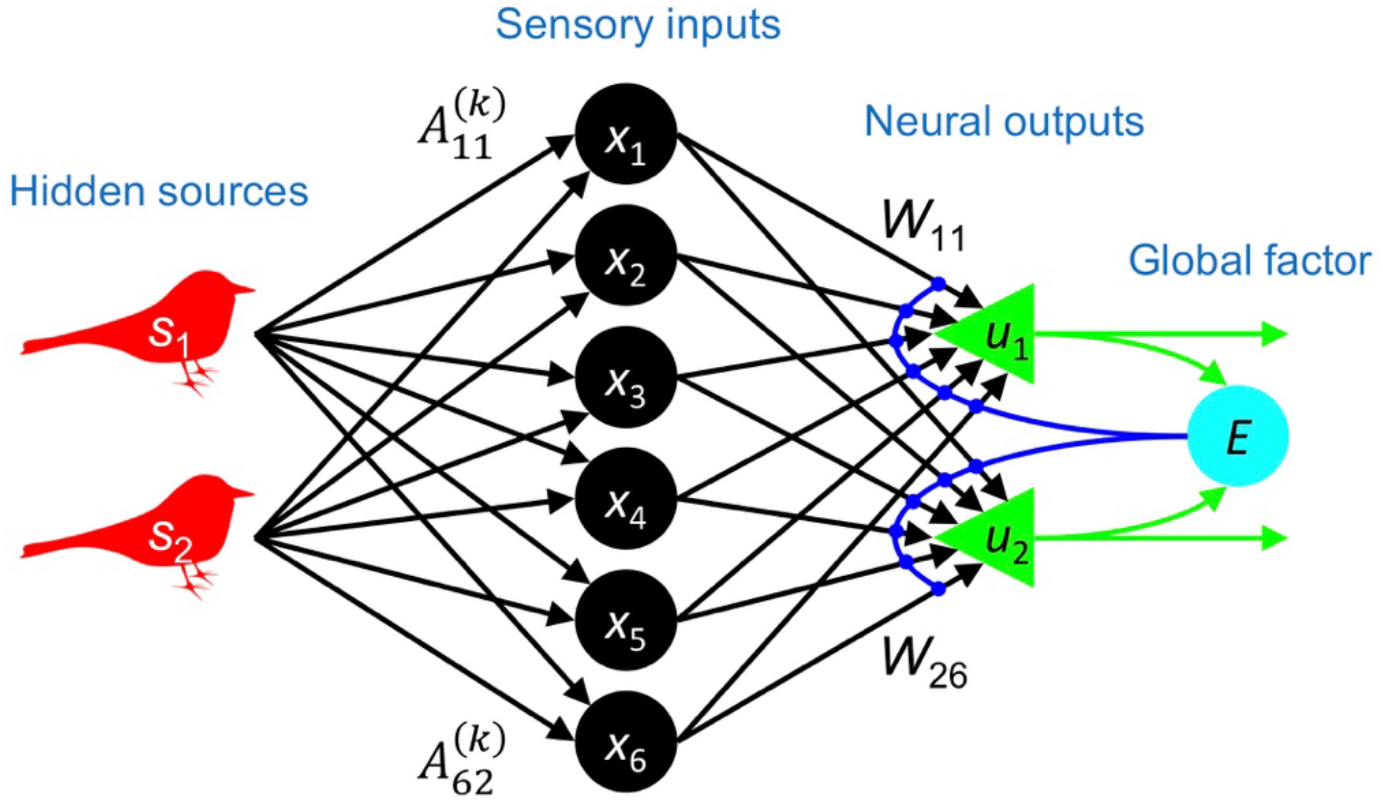
Model setup for multi-context BSS task. In this model, 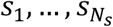 are hidden sources (e.g., birdsongs); 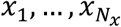 are sensory inputs that an agent receives; 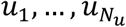 are neural outputs; 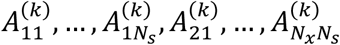 are elements of the *k*-th-context mixing matrix; 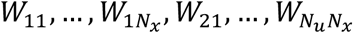 are synaptic strengths; and *E* is a scalar global factor that mediates synaptic plasticity. Synaptic strengths are adjusted to perform multi-context BSS by the EGHR.

Here *A*^(*k*)^ is the *N_x_* × *N_s_* mixing matrix for the *k*-th context that defines the magnitude of inputs, when each source generates a signal. To ensure that each *A*^(*k*)^ represents a different context and that each context has an ICA solution, column vectors of a block matrix (*A*^(1)^, *A*^(2)^,…,*A*^(*C*)^) are supposed to be linearly independent of each other. We design the task such that these contexts appear sequentially or randomly. The neural outputs are expressed as sums of inputs weighted by an *N_u_* × *N_x_* synaptic strength matrix *W*, and calculated by:

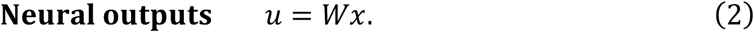

It is well known that when a presynaptic neuron (*x_j_*) and a postsynaptic neuron (*u_i_*) fire together, Hebbian plasticity occurs, and the synaptic connection from *x_j_* to *u_i_*, denoted by *W_ij_*, is strengthened (Hebb, 1949; Bliss, Lomo, 1973). Because this constitutes associative learning, correlations between 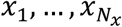 and *u_i_* are usually enhanced; thereby, correlations among neural outputs also increase. This is opposite to separation of signals (i.e., BSS) because each neural output is expected to encode a specific source to perform BSS. To separate signals, we introduce a global scalar factor (i.e., a third factor) given by the sum of nonlinearly-summed output units (*Isomura, Toyoizumi, 2016*):

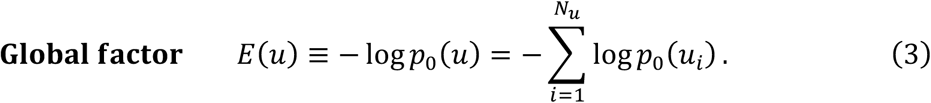

Here *p*_0_(*u*) is the prior distribution that the agent expects the hidden sources to follow; e.g., when *p*_0_(*u*) is a Laplace distribution of mean zero and unit variance, 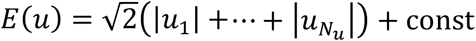. We suppose that this global factor modulates Hebbian plasticity. Recent experimental studies have reported that synaptic plasticity can be modulated by various neuromodulators (*Reynolds et al., 2001; Zhang et al., 2009; Salgado et al., 2012; Yagishita et al., 2014; Johansen et al., 2014*), GABAergic inputs (*Paille et al., 2013; Hayama et al., 2013*), or glial factors (*Ben Achour, Pascual, 2010*). Possible neurobiological implementations of the global factor are further discussed in the Discussion section. Overall, the synaptic strength matrix *W* is updated by the EGHR in the following way:

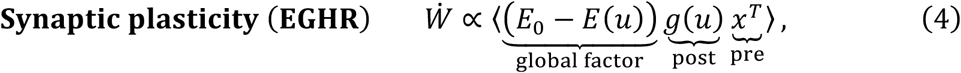

where *Ẇ* is the derivative of *W* with respect to time, 〈·〉 is the expectation over the input distribution, and *g*(*u*) ≡ d*E*(*u*) / d*u* is a non-linear function usually associated with a nonlinear activation function. A constant *E*_0_ scales the neural outputs; the output scale becomes equivalent to the source scale when *E*_0_ = E〈–log *p*_0_ (*s*)〉 + 1. In short, the EGHR constitutes a Hebbian learning rule when the global factor is smaller than the threshold (*E*(*u*) < *E*_0_), otherwise (*E*(*u*) > *E*_0_), it becomes an anti-Hebbian rule. This mechanism makes output neurons independent from each other. The detailed derivation and theoretical proofs of the EGHR have been described in previous reports (*Isomura, Toyoizumi, 2016; Isomura, Toyoizumi, 2018*).

### Memory capacity of the EGHR

First, we analytically show the memory capacity of a neural network established by the EGHR. As the number of contexts increases, larger dimensions of inputs are needed to retain information pertaining to past contexts in the neural network. For simplicity, let us suppose that *N_u_* = *N_s_*. Because the network represents a linear inverse model of the generative processes, the goal of the multi-context BSS is generally given by:

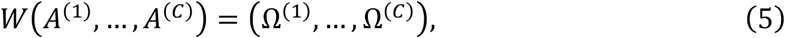

where Ω^(*k*)^ is an *N_u_* × *N_s_* matrix equivalent to the identity matrix, up to permutations and sign-flips. This is because the success of BSS is defined by one-by-one mapping from sources to outputs. Thus, if and only if the pseudo inverse matrix of a set of mixing matrices (*A*^(1)^,…, *A*^(*C*)^) is a full-rank, the multi-context BSS is successful. Hence, we find that the following conditions are necessary to achieve the multi-context BSS for a generic (*A*^(1)^,…, *A*^(*C*)^): (1) the input dimension needs to be equal to or larger than the number of contexts times the number of sources, *N_x_* ≥ *CN_s_*; and (2) the output dimension needs to be equal to or larger than the source dimension, *N_u_* ≥ *N_s_*. Note that the neural network learns the information representation compressing the sensory inputs because we consider much greater input than output dimensions.

The memory capacity of the EGHR was empirically confirmed by numerical simulations (Fig. 2). Here, we supposed that two contexts generated inputs alternately. In each context, six-dimensional inputs were generated from two-dimensional sources with different mixing weights, as denoted by *A*^(1)^ and *A*^(2)^ (see top and middle rows in Fig. 2A). A neural network consisting of six input and two output neurons received the inputs and changed its synaptic strengths through the EGHR (i.e., training). After training, each neural output came to selectively respond to (i.e., encode) one of the two sources (bottom row in Fig. 2A). Thus, the network achieved separation of the sensory inputs into their sources without being taught the mixing weights (i.e., BSS).

**Figure 2.**
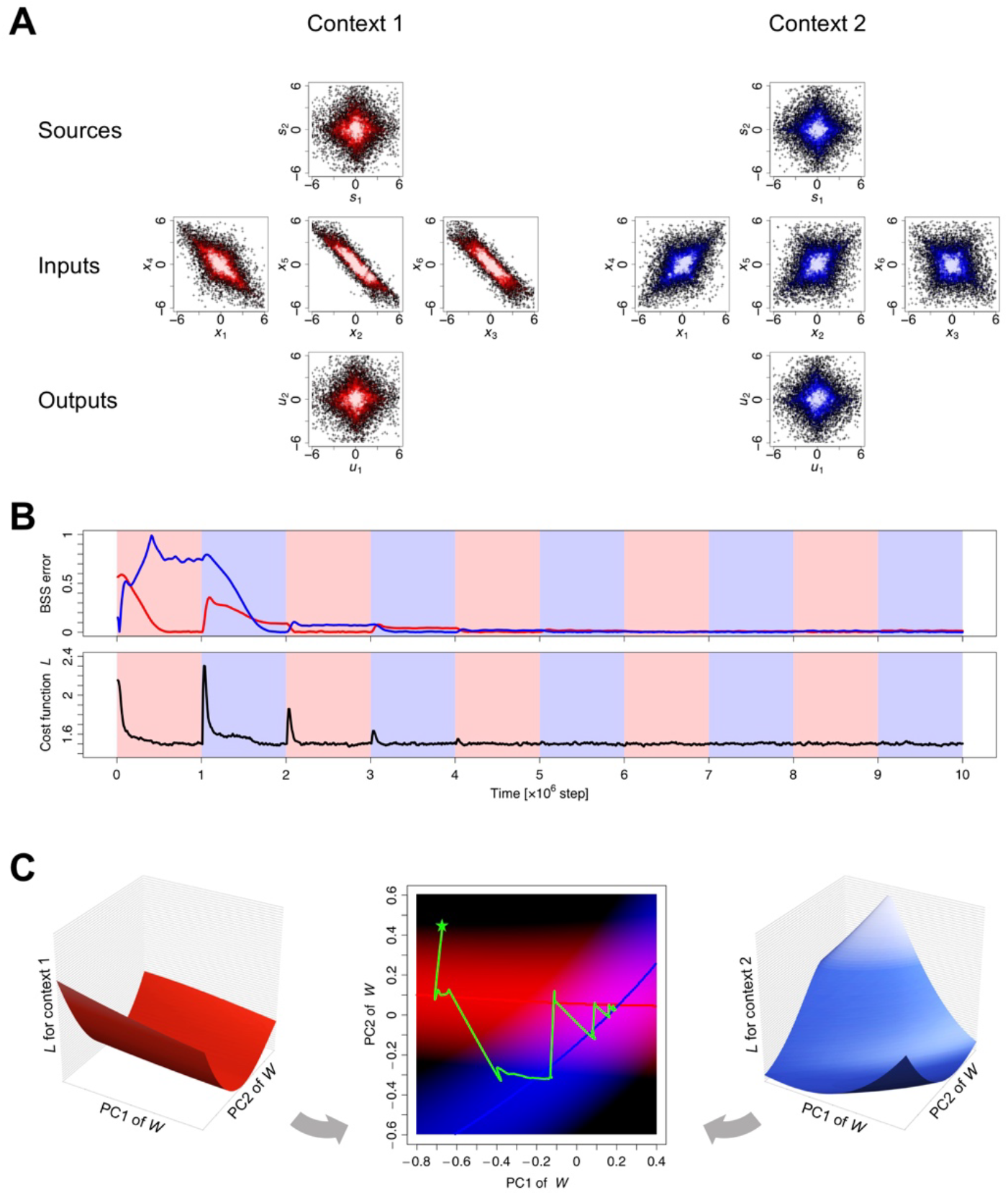
Results of multi-context BSS. (**A**) Distributions of sources, inputs, and outputs for context 1 and 2. (**B**) Trajectories of BSS error (top) and cost function (bottom). (**C**) Visualization of null spaces. The panel illustrates the shapes of the cost function under each context (left and right) and the trajectory of synaptic strengths (*W*) projected in a subspace spanned by the first (PC1) and second (PC2) principal components (center). The trajectory is determined by the gradient of either cost function, depending on the context. On the PC1-PC2 plane, null spaces are illustrated as nullclines; red and blue curves are nullclines for contexts 1 and 2, respectively. Low cost areas (i.e., valleys of the cost functions) are highlighted by red or blue shading. The synaptic strength matrix starts from a random initial state (star mark), shifts to the nullcline of context 1 or 2, and eventually converges to the cross point of the two nullclines, where the synaptic strengths perform the BSS for both contexts. Each source was randomly generated by the unit Laplace distribution. A learning rate of *η* = 4 × 10^−6^ was used. The MATLAB source code for this simulation is appended as Supplementary Source Codes.

Crucially, the neural network was able to retain the information learnt for all past contexts if provided with sufficiently redundant sensory inputs. This property is illustrated by the trajectories of the BSS error and EGHR cost function in Fig. 2B. We defined the BSS error for context *k* as the ratio of first to second maximum absolute values averaged for every row and column of matrix *K*^(*k*)^ ≡ *WA*^(*k*)^, which expresses the mapping from sources to outputs (equivalent to the covariance matrix between hidden sources and neural outputs *K*^(*k*)^ = Cov(*u*, *s*)). This definition was made to ensure that it took zero if and only if one source mapped onto one output, and *vice versa*; otherwise it took a positive value less than one. Moreover, the cost function of the EGHR was defined as the expectation of the square of the global factor: *L* ≡ 〈(*E*_0_ – *E*(*u*))^2^/2〉. Context 1 (red in Fig. 2A) was provided in the first session. Since synaptic strengths started from a random initial state, the BSS error at the beginning of the first session was large; then, the network learned an optimal set of synaptic strengths, and the error became zero. Notably, the EGHR was defined to reduce the EGHR cost function by gradient descent updates. However, when context 2 (blue in Fig. 2A) was provided for the first time in the second session, the EGHR cost function transiently increased because it needed to learn the new mixing matrix. An important point was revealed at the first step of the third session, in which context 1 was provided again. The BSS error was significantly smaller than that in the first session and close to zero from the beginning of this session, indicating that the network retained synaptic strengths that were optimized for context 1 even after learning context 2. After several iterations, the BSS error for both contexts converged to zero. The success of learning was also confirmed by the trajectory of the EGHR cost function that also converged to the minimum value.

These results show that an undercomplete EGHR increased the speed of re-adaptation to previously experienced contexts, suggesting that memory of past experiences was preserved within the network. Moreover, the network learned the optimal set of synaptic strengths that entertained both contexts after several iterations. A key feature for this ability is the “null space” in the synaptic strength matrix. Since only 2 × 2 dimensions were required to express a mapping from two-dimensional sources to two-dimensional outputs in one context, the synaptic strength matrix comprised eight (2 × 6 – 2 × 2)-dimensional degrees of freedom. This freedom spanned a null space in which synaptic strengths were equally optimized with a zero BSS error. Similarly, when two different contexts were considered, four-dimensional degrees of freedom remained, as an overlap between the two eight-dimensional null spaces. To visualize such a null space, we projected synaptic strengths onto a subspace spanned by the first (PC1) and second (PC2) principal components of the trajectory of synaptic strengths. On this PC1-PC2 plane, a null space was illustrated as a nullcline. Since the dynamics of synaptic strengths were determined to go down the slope of a cost function for either context 1 or 2, synaptic strengths were started from a random initial state and reached the nullcline of either context 1 or 2, in turn. Crucially, this trajectory converged to the cross point of two nullclines, where the synaptic strengths entertained both contexts. Because of this, the BSS error reached zero after iterative training, i.e., an ICA solution that was optimized for both contexts was found.

Furthermore, we examined the multi-context BSS by the EGHR using a large number of contexts (Fig. 3). Our agent received redundant (2000-dimensional) sensory inputs, comprising 100 sets (contexts) of mixtures of ten hidden sources (1000 sources in total), that were generated as products of the context-dependent mixing matrix and sources. Ten outputs neurons learned to infer each source from their mixtures by updating synaptic strengths through the EGHR. After training, we found that they successfully represented the ten sources for every context, without further updating synaptic strengths, as illustrated by the reduction of the BSS error for all 100 contexts (Fig. 3A) and the convergence of the covariance between sources and outputs to a diagonal matrix (up to permutations and sign-flips) (Fig. 3B). This was because synaptic strengths had sufficient capacity and were formed to express the inverse of the concatenated mixing matrices from all contexts, which was further confirmed by the convergence of the synaptic strength matrix in the null space (Fig. 3C).

**Figure 3.**
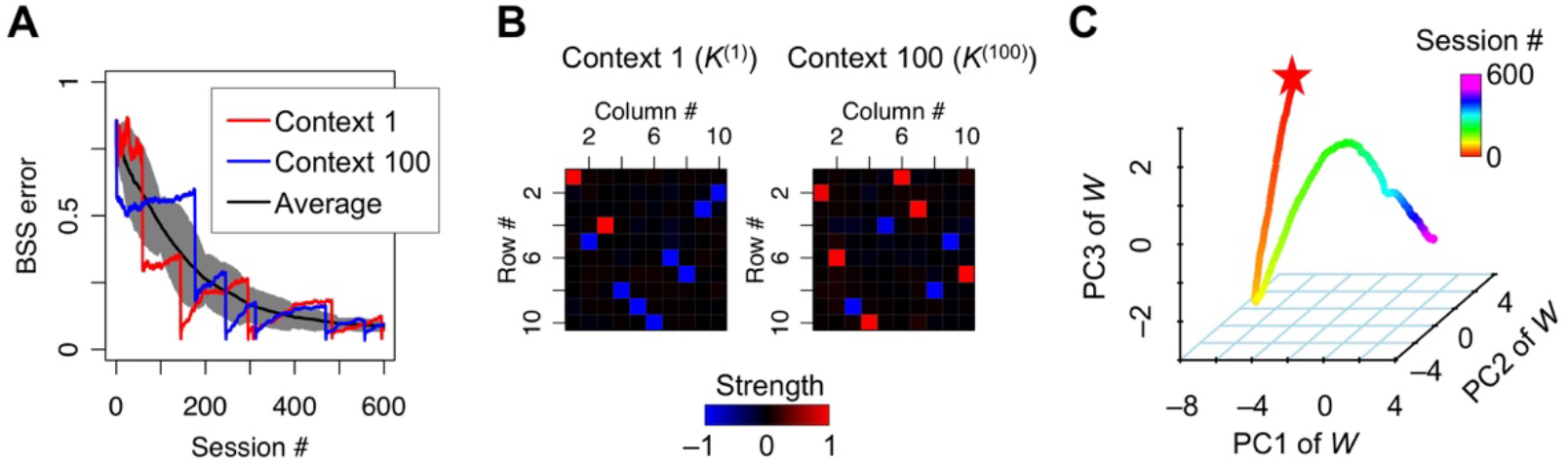
BSS with large number of contexts. One of 100 different contexts was randomly selected for each session. Each session contained *T* = 10^5^ time steps and the training continued 600 sessions. In each session, the 2000-dimensional sensory inputs (*x*) were generated from ten-dimensional hidden sources (*s*), which independently followed the unit Laplace distribution, through a context-dependent random mixing matrix *A*^(*k*)^. The neural network consisting of ten-dimensional neural outputs (*u*) was trained with a learning rate of *η* = 1 × 10^−5^. (**A**) Trajectories of BSS error for context 1 and 100 and the average BSS error over contexts 1 to 100. The shaded area shows the standard deviation. (**B**) Mappings from ten sources to ten outputs in contexts 1 and 100 after training. Elements of matrix *K*^(*k*)^ = *W A*^(*k*)^ with *k* = 1 and 100 are illustrated by the heat map. Only one element in each row and column takes ±1, indicating the one-to-one mapping from sources to outputs, i.e., the success of multi-context BSS. (**C**) The dynamics of synaptic strength matrix *W* projected in the three-dimensional space spanned by the first to third principal components (PC1 to PC3). The matrix starts from a random initial point (star mark) and converges to the null space, in which synaptic strengths are optimized for all trained contexts. The C code for this simulation is appended as Supplementary Source Codes.

### BSS in constantly time-varying environments

In the previous section, we described a general condition for the neural network to achieve the multi-context BSS. In special cases, where the mixing matrices in each context have common features, the neural network can perform BSS beyond the capacity mentioned above. Here we show that when contexts are generated from a low-dimensional subspace of mixing matrices and thereby are dependent on each other, the EGHR can find the common features and use them to perform the multi-context BSS.

As a corollary of the property of the EGHR when provided with redundant inputs, the EGHR can perform the BSS even when the mixing matrix changes constantly as a function of time (Fig. 4A). Without loss of generality, a time-dependent mixing matrix is expressed by the sum of time-invariant and time-variant components, as follows:

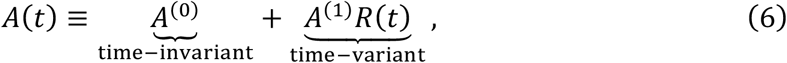

where *A*^(0)^ is a constant matrix with the same size as *A*(*t*), *A*^(1)^ is a constant vertically-long rectangular (or square) matrix, and *R*(*t*) is a matrix composed of either smoothly or discontinuously changing functions in time. Each component of *R*(*t*) is supposed to on average slowly change, i.e., their time-derivatives are typically much smaller in magnitude than those of *s*(*t*). This condition is required to distinguish whether changes in inputs are caused by changes in the mixing matrix *A*(*t*) or the hidden sources *s*(*t*). Formally, *A*(*t*) expresses infinite contexts along the trajectory of *R*(*t*). This is a more complicated setup than the standard BSS in the sense that both sources and the mixing matrix change in time. Nonetheless, the EGHR can achieve BSS for all contexts if a solution of the synaptic strength matrix that satisfies *W*(*A*^(0)^, *A*^(1)^) = (*Ω*, *O*) exists. Here, Ω represents the identity matrix up to permutations and sign-flips. Such a solution generally exists if *A*^(0)^ is a full rank matrix, and *N_x_* is greater than the sum of the source dimension and the row dimension of *R*(*t*). The above condition means that the network performs BSS based on the time-invariant features *A*^(0)^ of the mixing matrix, while neglecting the time-varying features *A*^(1)^*R*^(*t*)^. This can be viewed as a way to compress high-dimensional data. This is distinct from the standard dimensionality reduction approach by PCA, which would preferentially extract the time-variant features due to their extra variances.

**Figure 4.**
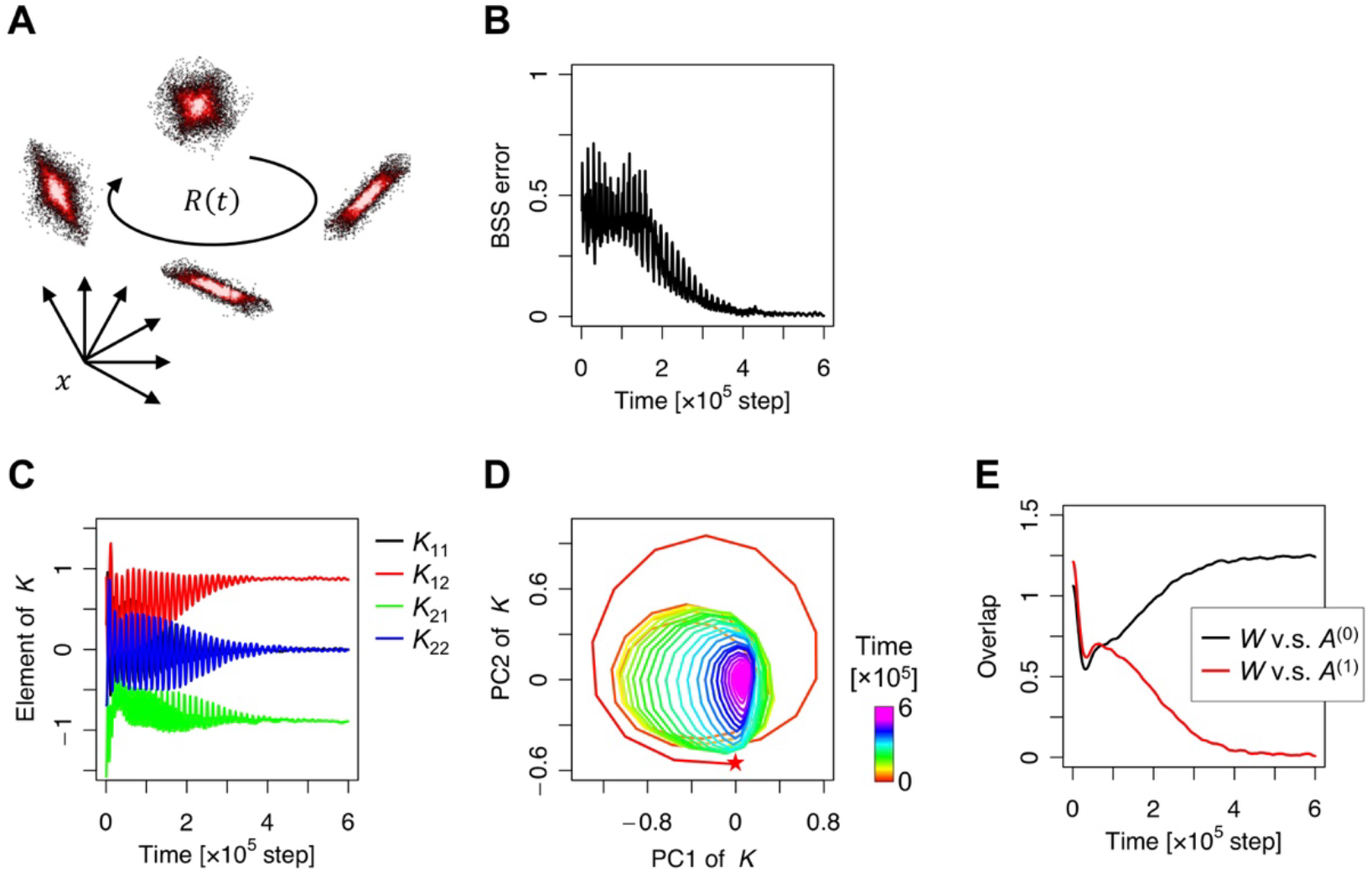
BSS with time-varying mixing matrix. (**A**) Schematic image of sensory inputs generated from two sources through time-varying mixing matrix *A*(*t*). The mixing matrix is controlled by the low-dimensional rotation matrix *R*(*t*). (**B**) Trajectory of BSS error. (**C**) Trajectories of mapping weights from sources to outputs, i.e., matrix *K* = *W A*(*t*). (**D**) Dynamics of matrix *K* projected on the first two-dimensional PCA subspace of *K*’s trajectory over training. The matrix starts from a random initial point (star mark) and follows a spiral trajectory as it converges to a subspace in which synaptic matrix *W* is perpendicular to the time-varying component *A*^(1)^. (**E**) Overlap of synaptic matrix *W* with time-invariant component *A*^(0)^ and time-variant component *A*^(1)^. The overlap between two matrices was defined by the Frobenius norm of their product, i.e., 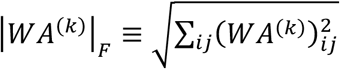. Each source was randomly generated by the unit Laplace distribution. The learning rate of *η* = 1 × 10^−5^ was used. The C code for this simulation is appended as Supplementary Source Codes.

In the simulation, we supposed *R*(*t*) to be a two-dimensional rotation matrix, *R*(*t*) = (cos *ωt*, − sin *ωt*; sin *ωt*, cos *ωt*), with an angular frequency of 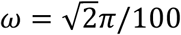. The simulation showed a reduction in the BSS error (Fig. 4B). At the same time, *K* = *WA* converged to the identity matrix up to permutations and sign-flips, *K*→ (0,1; −1,0) in this case, although *A* continuously changed in time (Figs. 4C and 4D). As illustrated in Fig. 4E, the synaptic matrix *W* became perpendicular to the time-varying features *A*^(1)^ (i.e., *WA*^(1)^ = *O*), by a monotonic reduction of the overlap between *W* and *A*^(1)^ (defined by the Frobenius norm of their product). After training, the overlap converged to zero. Hence, synaptic strengths were optimized regardless of *R*(*t*) at this solution, which enabled the network to perform BSS with a virtually infinite number of contexts.

Next, we demonstrated the utility of the EGHR, when supplied with redundant inputs, by using natural birdsongs and a time-variant mixing matrix that expressed a natural contextual change. Figure 5 illustrates the BSS task of two birdsongs when birds moved around the agent; thereby, the mixing matrix changed in time according to the positions of the birds. Please see also Supplementary Movie S1 for a comparison of the neural outputs before and after training. To obtain time-independent features, we assumed that the two birds moved around in non-overlapping areas. For simplicity, we also assumed that the two birds moved around at different heights. The agent received mixtures of the two birdsongs through six microphones with different direction preferences regarding the z-axis (i.e., time-invariant features) and the x-and y-axes (i.e., time-variant features). By tuning synaptic strengths by the EGHR, neural outputs were established to infer each birdsong, while the mixing matrix changed continuously. Crucially, after training, the mapping from the sources to the outputs (*K* = *W A*) became constant with time, although matrix *A* was time-dependent. More precisely, the EGHR found a representation where *W* satisfied *W*(*A*^(0)^, *A*^(1)^) = (Ω, *O*). Hence, neural outputs could separate the two birdsongs, although the amplitudes of the songs recorded by the microphones continuously changed depending on the positions of birds.

**Figure 5.**
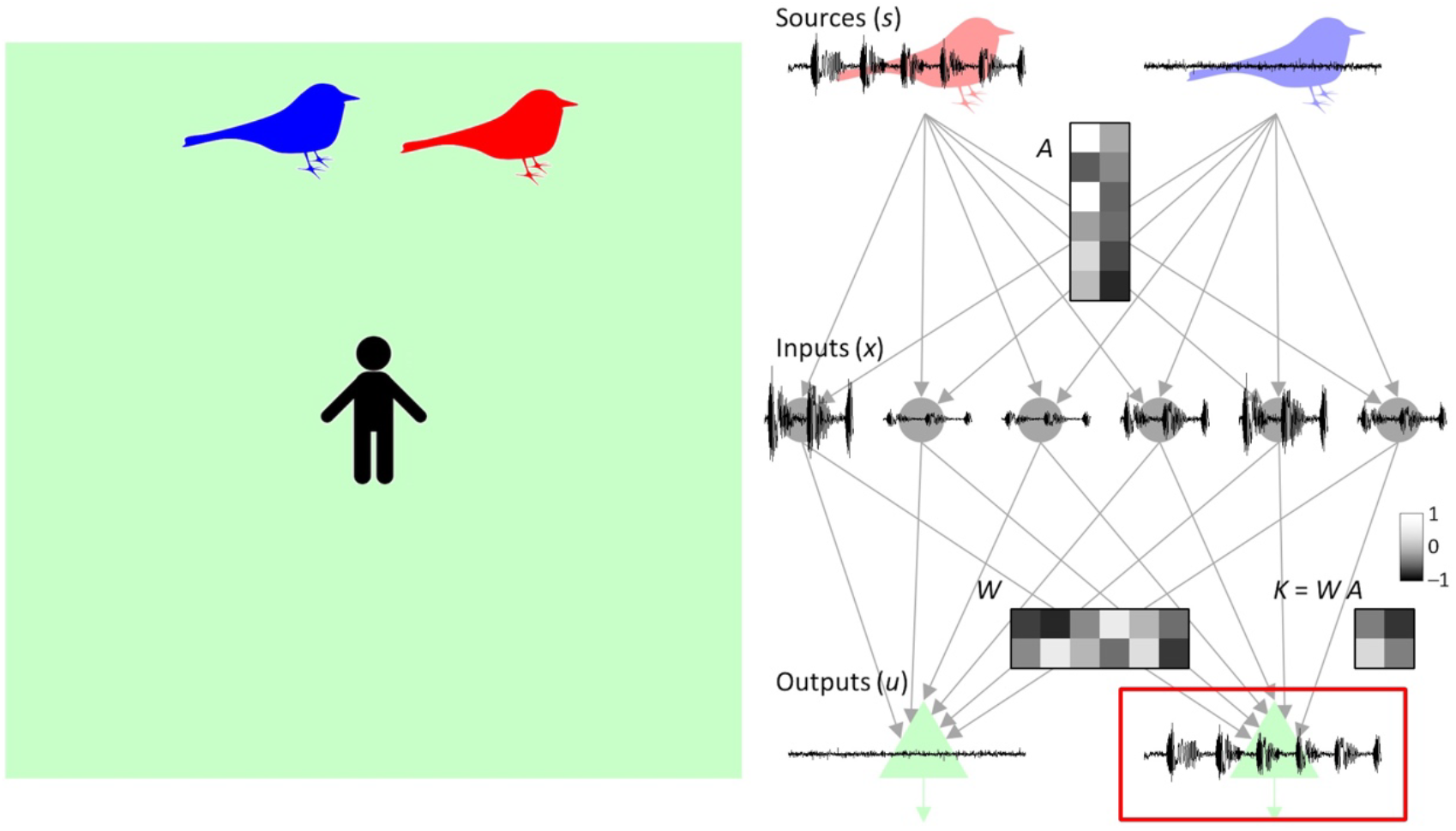
BSS of birdsongs when two birds move around the agent. A snapshot of the simulation overview movie after training is shown. Songs (or sources) generated by two birds *s*_1_, *s*_2_ (right top) are mixed with a time-varying mixing matrix *A,* resulting in six-dimensional sensory inputs *x*_1_,…, *x*_6_ (right middle). The mixed signals correspond to the recording through six microphones with different preferences. The neural network converts the six inputs into two neural outputs *u*_1_, *u*_2_ (right bottom) using the synaptic strength matrix *W*. The synaptic updates by the EGHR enable the outputs to encode each birdsong. Matrix *K* = *W A* represents the mapping from sources to outputs. See Methods for the detailed simulation setup.

### Generalization for inexperienced environments

Finally, we examined the generalization capability of the multi-context BSS by the EGHR using natural birdsongs. For the sake of simplicity, we reduced Eq. (6) by considering *R*(*t*) that changes discontinuously at the beginning of each session but otherwise is constant. Specifically, we considered the mixing matrix

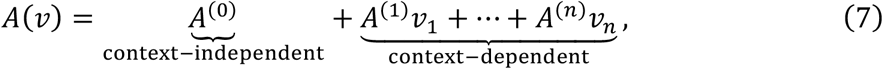

written using context-independent matrix *A*^(0)^, context-dependent matrices {*A*^(1)^,…,*A*^(*n*)^}, and context vector *v* ≡ (*v*_1_,…, *v_n_*) that discontinuously changes at the beginning of a new session. The first term in the right-hand side of Eq. (7) corresponds to the context-independent (i.e., constant) component, which should be a full-rank matrix to provide an ICA solution. Similarly to the case with the continuously time-varying mixing matrix, the EGHR can establish synaptic matrix *W* that expresses the pseudo inverse of *A*^(0)^ up to permutations and sign-flips, while keeping *W* perpendicular to *A*^(1)^,…,*A*^(*C*)^, i.e., *W*(*A*^(0)^, *A*^(1)^,…,*A*^(*C*)^) = (Ω, *O*,…, *O*). Notably, the EGHR can establish such *W* by using only a handful samples of *v* out of combinatorially many possibilities. This is because the mappings from sources to inputs are restricted to be a linear transformation and thereby observations with the polynomial (probably quadratic) order number of contexts can identify the mapping for all contexts. This property is particularly useful when *v* is high dimensional.

In this demonstration, ten sets (contexts) of mixtures of ten birdsongs were introduced to our agent, with redundant sensory inputs composed of 100 mixed sound waves (Fig. 6A). Those contexts were defined by random mixing matrices *A*^(0)^, *A*^(1)^,…, *A*^(4)^. We trained the network using only 10 contexts: *v* = (1,0,0,0), (½,½,0,0), (0,1,0,0), (0,½,½,0), (0,0,1,0), (0,0,½,½), (0,0,0,1), (½,0,0,½), (½,0,½,0), (0,½,0,½). At the beginning of each session, *v* was randomly selected from the above ten vectors, which provided a discrete random transition among 10 contexts. Ten output neurons learned to infer each birdsong from their mixtures, by updating synaptic strengths through the EGHR. After training, they successfully represented the ten birdsongs without further updating synaptic strengths. Crucially, the network could perform BSS even in an inexperienced context (for example, in *v* = (¼,¼,¼,¼); the most right column in Fig. 6A). This speaks to the generalization of the multi-context BSS for unseen test contexts.

**Figure 6.**
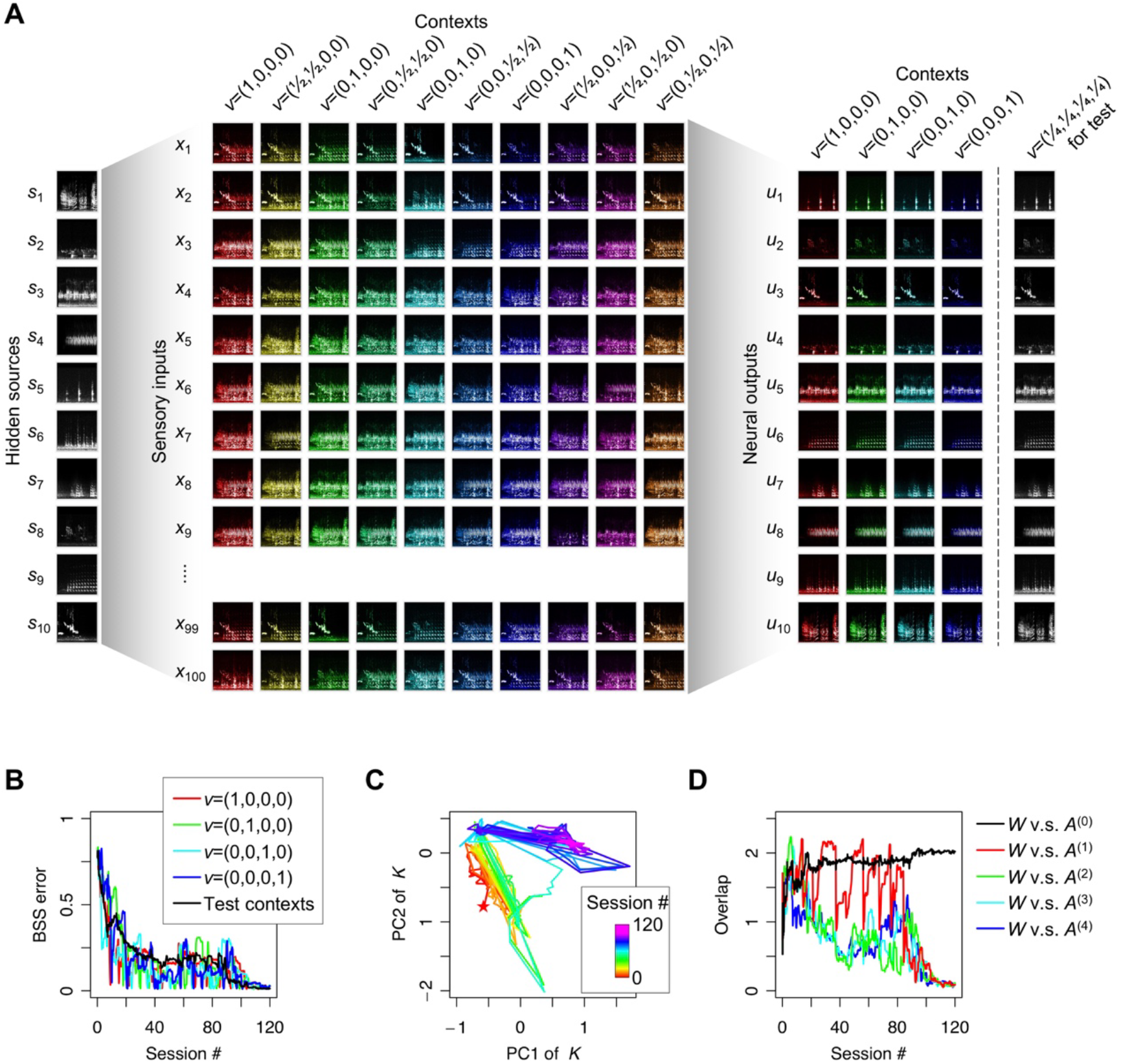
Generalization of multi-context BSS. (**A**) Spectrograms of ten hidden sources (birdsongs), 100 sensory inputs, and ten neural outputs. Each context is illustrated in a different color. (**B**) Trajectories of the BSS error with four trained contexts and the average BSS error over 20 inexperienced test contexts, created using randomly sampled *v*. (**C**) Dynamics of matrix *K* projected on the first two-dimensional PCA subspace. The matrix starts from a random initial point (star mark) and converges to a fixed point, at which the synaptic matrix entertains every trained context. (**D**) Overlap of synaptic matrix *W* with context-independent component *A*^(0)^ and context-dependent components *A*^(1)^,…,*A*^(4)^). Overlap between two matrices is defined by |*WA*^(*k*)^|_*F*_, as described in Fig. 4. See Methods for the detailed simulation setup.

Next, we quantitatively showed that, as learning progresses, the BSS error for test contexts (defined using 20 randomly sampled *v* from a uniform distribution [0,1] that were inexperienced in the training), as well as trained contexts, decreased (Fig. 6B). Regardless of the given context, matrix *K* converged to a constant matrix that was the same as the identity matrix up to permutations and sign-flips, as illustrated by the convergence of principal components of *K* into a fixed point representing the context-independent null space (Fig. 6C).

This convergence was further validated by plotting the trajectories of the overlaps between *W* and *A* components (Fig. 6D). While the overlap between *W* and *A*^(0)^ increased as learning progressed, the overlap with context-dependent components (*A*^(1)^,…,*A*^(4)^) decreased and converged to zero, showing that *W* became perpendicular to *A*^(1)^,…,*A*^(4)^ by the EGHR. Hence, this learnt network could perform BSS with *A*(*v*) determined by arbitrary *v* in the four-dimensional space, without further synaptic updating or transient error, while the network was trained only with 10 contexts. Those results highlight the significant generalization capability of the neural network established by the EGHR and the robustness against inexperienced environments for performing BSS.

## Discussion

While a real environment comprises several different contexts, humans and animals retain the experience of past contexts to perform well when they find themselves in the same context in the future. This ability is known as conservation of learning or cognitive flexibility (*Mante et al, 2013; Dajani, Uddin, 2015*). Although analogous learning is likely to happen during BSS, the conventional biological BSS algorithms (*Foldiak, 1990; Linsker, 1997*) must forget the memory of past contexts to learn a new one. Thereby, when the agent subsequently encounters a previously experienced context, it needs to relearn it from the very beginning. We overcame this limitation by using the described algorithm, the EGHR. The crucial property of the EGHR is that when the number of inputs is larger than the number of sources, the synaptic matrix contains a null space in which synaptic strengths are equally optimized for performing BSS. Hence, with sufficiently redundant inputs, the EGHR can make the synaptic matrix optimal for every experienced context. This is an ability that the conventional biologically plausible BSS algorithms do not have, due to the constraint that the number of inputs and outputs must be equal; however, this ability is crucial for animals to perceive and adapt to dynamically changing multi-context environments. It is also crucial for animals to generalize past learning to inexperienced contexts. We also found that, if there is a common feature shared across the training contexts, the EGHR can extract it and generalize the BSS result to inexperienced test contexts. This speaks to a generalization capability and transfer learning, i.e., this result implies prevention of overfitting to a specific context; alternatively, one might see this as an extraction of a general concept across the contexts. Therefore, we argue that the EGHR is a good candidate model for describing the neural mechanism of conservation of learning or cognitive flexibility for BSS.

Moreover, the process of extracting hidden sources in the multi-context BSS setup can be seen as a novel concept of dimensionality reduction (*Cunningham, Ghahramani, 2015*). If the dimensions of input are greater than the product of the number of sources and the number of contexts, the EGHR can extract the low-dimensional sources (up to context-dependent permutations and sign-flips), while filtering out a large number of context-dependent signals induced by changes in the mixing matrix. ICA algorithms for multi-context BSS (*Amari et al., 2000; Lee, Lewicki et al., 2000; Hirayama et al., 2015*) and undercomplete ICA for compressing data dimensionality (*Porill and Stone, 1997; Isomura and Toyoizumi, 2016; Isomura and Toyoizumi, 2018*) have been separately developed. Nevertheless, to our knowledge, our study is the first to attempt a dimensionality reduction in the multi-context BSS setup. This method is particularly powerful when a common feature is shared across the contexts, because the EGHR can make each neuron encode an identical source across all the contexts. Our results are different from those obtained using standard dimensionality reduction approaches by PCA (*Pearson, 1901; Oja, 1989*), because PCA is used for extracting subspaces of high-variance principal components and hence would preferentially extract the context-dependent varying features, given that each source has the same variance. Therefore, our study proposes an attractive use of the EGHR for dimensionality reduction.

We demonstrated that a neural network learns to distinguish individual birdsongs from their superposition. Young songbirds learn songs by mimicking adult birds’ songs (*Tchernichovski et al., 2001; Woolley, 2012; Lipkind et al., 2013; Lipkind et al., 2017*). A study reported that neurons in songbirds’ higher auditory cortex exhibit a teacher specific activity (*Yanagihara, Yazaki-Sugiyama, 2016*). One can imagine those neurons correspond to the expectation of hidden sources (*u*), as considered in this study. Importantly, the natural environment that young songbirds encounter is dynamic, as we considered in Fig. 5. Therefore, the conventional BSS setup, which assumes a static environment or context, is not suitable for explaining this problem. It is interesting to consider that young songbirds might employ some computational mechanism similar to the EGHR to distinguish a teacher song from other songs in a dynamically changing environment.

It is worth noting that the application of standard ICA algorithms to high-pass filtered inputs cannot solve the multi-context BSS problem. This is because context-dependent changes in the mixing matrix not only change the means of the inputs, which can be removed by high-pass filtering, but also change the gain of how each source fluctuations are propagated to input fluctuations. Hence, the difference in contexts cannot be expressed as a linear ICA problem after high-pass input filtering. Therefore, selective extraction of context-invariant features is an advantage of the EGHR. Moreover, if provided with redundant input, the EGHR can solve multi-context BSS even if the context changes continuously in time, as we demonstrated in Figs. 4, 5.

Biological neural networks implement an EGHR-like learning rule. The main ingredients of the EGHR are Hebbian plasticity and the third scalar factor that modulates it. Hebbian plasticity occurs in the brain depending on the activity level (*Bliss, Lomo, 1973; Dudek and Bear, 1992*), spike timings (*Markram et al., 1997; Bi, Poo, 1998; Zhang et al. 1998; Feldman, 2012*), or burst timings (*Butts et al. 2007*) of pre-and post-synaptic neurons. In contrast, the third scalar factor can modify the learning rate and even invert Hebbian to anti-Hebbian plasticity (*Paille et al., 2013*), similarly to what we propose for the EGHR. In general, such a modulation forms the basis of a three-factor learning rule, a concept that has recently received attention, see (*Pawlak et al., 2010; Frémaux, Gerstner, 2016; Kuśmierz et al., 2017*) for reviews, and is supported by experiments on various neuromodulators and neurotransmitters, such as dopamine (*Reynolds et al., 2001; Zhang et al., 2009; Yagishita et al., 2014*), noradrenaline (*Salgado et al., 2012; Johansen et al., 2014*), and GABAergic input (*Paille et al., 2013; Hayama et al., 2013*), as well as glial factors (*Ben Achour, Pascual, 2010*). Those third factors may encode reward (*Izhikevich, 2007; Florian, 2007; Legenstein et al., 2008; Urbanczik, Senn, 2009; Frémaux et al., 2010*), likelihood (*Brea et al., 2013*), novelty/surprise (*Rezende, Gerstner, 2014*), or error from a prior belief (*Isomura, Toyoizumi, 2016; Isomura, Toyoizumi, 2018*). Importantly, the EGHR only requires such local signals, which neurons can access via their synaptic connections. Furthermore, a study using *in vitro* neural networks suggested that neurons perform simple BSS using a plasticity rule that is different from the most basic form of Hebbian plasticity, by which synaptic strengths are updated purely as a product of pre-and postsynaptic activity (*Isomura et al., 2015*). A candidate implementation of the EGHR can be made for cortical pyramidal cells and inhibitory neurons; the former constituting the EGHR output neurons and encoding the expectations of hidden sources, and the latter constituting the third scalar factor and calculating the nonlinear sum of activity in surrounding pyramidal cells. This view is consistent with the circuit structure reported for the visual cortex (*Hofer et al., 2011; Harris, Mrsic-Flogel, 2013*). These empirical evidences support the biological plausibility of the EGHR as a candidate model of neuronal BSS.

A local computation of the EGHR is highly desirable for neuromorphic engineering (*Chicca et al., 2014; Merolla et al., 2014; Neftci, 2018*). The EGHR updates synapses by a simple product of pre-and postsynaptic neurons’ activity and a global scalar factor. Because of this, less information transfer between neurons is required, compared to conventional ICA methods that require non-local information (*Bell, Sejnowski, 1995; Bell, Sejnowski, 1997; Amari et al., 1996*), all-to-all plastic lateral inhibition between output neurons (*Foldiak, 1990; Linsker, 1997*), or an additional processing step for decorrelation (*Hyvarinen, Oja, 1997*). The simplicity of the EGHR is a great advantage when implemented in a neuromorphic chip because it can reduce the space for wiring and the energy consumption. Furthermore, unlike the conventional ICA algorithms that assume an equal number of input and output neurons, a neuromorphic chip that employs the EGHR with redundant inputs would perform BSS in multiple contexts, as allowed by the network memory capacity, without requiring readaptation. The generalization capability of the EGHR, as demonstrated in Fig. 6, is an additional benefit, as the EGHR captures the common features shared across training contexts to perform BSS in inexperienced test contexts.

Notably, although we considered the linear BSS problem in this study, multi-context BSS can be extended to a non-linear BSS, in which the inputs are generated through a non-linear mixture of sources (*Lappalainen, Honkela, 2000; Karhunen, 2001*). To solve this problem, a promising approach would be to use a linear neural network. A recent study showed that when the ratio of input-to-source dimensions and source number are large, a linear neural network can find an optimal linear encoder that separates the true sources through PCA and ICA, thus asymptotically achieving zero BSS error (*Isomura, Toyoizumi, 2018b*). Because both the asymptotic linearization and multi-context BSS by the EGHR are based on high-dimensional sensory inputs, it would be useful to combine them to solve the multi-context, non-linear BSS problem.

In summary, we demonstrated that the EGHR can retain memories of past contexts and, once the learning is achieved for every context, it can perform multi-context BSS without further updating synapses. Moreover, the EGHR can find common features shared across contexts, if present, and uses them to generalize the learning result to inexperienced contexts. Therefore, the EGHR will be useful for understanding the neural mechanisms of flexible inference and sensory representation under dynamically changing environments.

## Methods

### Model and learning rule

The neural network model and used learning rule (the EGHR) are described in the Results section.

### BSS solution

Suppose that *N_u_* = *N_s_*. We define a transform matrix *K*^(*k*)^ by

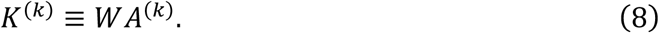

For *N_x_* ≥ *N_s_*, the ICA for context *k* is achieved when *K*^(*k*)^ is the identical matrix up to permutations and sign-flips. Hence, when column vectors of a block matrix (*A*^(1)^,…, *A*^(*C*)^) are independent of each other, if and only if (*A*^(1)^,…, *A*^(*C*)^) is a full-rank matrix, an ICA solution that separates all sources for context 1,…, *C* exists. Namely, *W* achieves the multi-context BSS when it satisfies

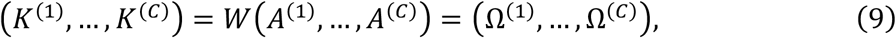

where Ω^(*k*)^ is an *N_u_* × *N_s_* matrix equivalent to the identity matrix up to permutations and sign-flips. Regarding the *i*-th row of matrix *K*^(*k*)^, as denoted by a row vector 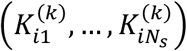, the achievement of ICA is justified when one element is one and the others are zero. Thus, there are many candidate sets of 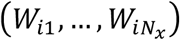 that can achieve ICA, because *N_x_* is larger than *N_s_*. Our numerical analyses showed that among these potential solutions, the one that is the nearest to the solution for the previous context is likely to be chosen. This can be understood as follows: when the network finds an ICA solution for all contexts, the error (i.e., cost function of the EGHR) is minimized including transient periods between two contexts; hence, according to gradient descent, synaptic strengths converge to such a solution as training progresses. Owing to this mechanism, the initial errors converge to zero when previously experienced environments are provided as stimuli.

### Simulation protocols

#### For figure 5

Two birdsongs were downloaded from Xeno-canto (https://www.xeno-canto.org/132149, https://www.xeno-canto.org/133054). Two hidden sources were created by trimming the first 60 s of these songs (with 4410-Hz time resolution) and normalizing them, to ensure each source sequence had zero mean and unit variance. During the training, the song sequences were repeated. To add stochasticity, a hidden source was defined by the sum of a song and a white-noise sequence generated by a Laplace distribution. The mixing matrix was defined by 6 × 2 random matrices, *A*^(0)^, *A*^(1)^, and a rotation matrix, *R*(*t*) ≡ (cos *ωt*, − sin *ωt*; sin *ωt*, cos *ωt*). The angular frequency *ω* was randomly set as −0.1*π*, 0, or 0.1*π* [rad/s], by following Poisson process with a transition probability of 1/8820. The training time and learning rate were defined by *T* = 4410 × 6000 [step] and *η* = 1 × 10^−7^.

#### For figure 6

Ten birdsongs were downloaded from Xeno-canto (the URLs are https://www.xeno-canto.org/****** where ****** was replaced with the following numbers: 27060, 64735, 67307, 110303, 121326, 121691, 126481, 132149, 133054, 133862). Ten hidden sources were created in the same manner as described above. The mixing matrix was defined by 100 × 10 random matrices *A*^(0)^, *A*^(1)^, *A*^(2)^, *A*^(3)^, *A*^(4)^, where *Â*^(1)^,…,*Â*^(4)^ were randomly generated and *A*^(0)^,…,*A*^(4)^ were defined by A(0) = (*Â*^(1)^ + *Â*^(2)^ + *Â*^(3)^ + *Â*^(4)^)/ 4 and *A*^(*k*)^ = *Â*^(*k*)^ = *A*^(0)^, for *k* = 1,…, 4. This treatment served to ensure that *A*^(1)^,…*A*^(4)^ do not involve common features across contexts. The training comprised 120 sessions, with each session continued for *T* = 4410 × 600 [step]. The context vector *v* randomly chose one of the following ten vectors, *v* = (1,0,0,0), (½,½,0,0), (0,1,0,0), (0,½,½,0), (0,0,1,0), (0,0,½,½), (0,0,0,1), (½,0,0,½), (½,0,½,0), (0,½,0,½), at the beginning of each session and maintained the value during the session. The learning rate was defined by *η* = 2 × 10^−7^. For the test, 20 randomly generated vectors were used, and their elements were randomly sampled from [0,1] and then normalized to satisfy *v*_1_ + *v*_2_ + *v*_3_ + *v*_4_ = 1.

## Acknowledgements

This work was supported by RIKEN Center for Brain Science (T.I. and T.T.), Brain/MINDS from AMED under Grant Number JP18dm020700 (T.T.), and JSPS KAKENHI Grant Number JP18H05432 (T.T.). The funders had no role in study design, data collection and analysis, decision to publish, or preparation of the manuscript.

## Author Contributions

Conceived the idea: T.I. and T.T. Performed the analyses: T.I. and T.T. Wrote the paper: T.I. and T.T.

